# Proteomics and Functional Analysis Revealed TaGSTU6/TaCBSX3 Enhances Wheat Resistance to Powdery Mildew

**DOI:** 10.1101/2022.01.31.478538

**Authors:** Qiao Wang, Jia Guo, Pengfei Jin, Guo Meng Ying, Jun Guo, Cheng Peng, Qiang Li, Baotong Wang

## Abstract

Wheat stripe rust and powdery mildew are important worldwide diseases of wheat (*Triticum aestivum*). The wheat cultivar Xingmin318 (XM318) is resistant to both wheat stripe rust and powdery mildew, which are caused by *Puccinia striiformis* f. sp. *tritici* (*Pst*) and *Blumeria graminis* f. sp. *tritici* (*Bgt*), respectively. To explore the molecular mechanisms of wheat defenses against *Pst* and *Bgt*, quantitative proteomic analyses of XM318 inoculated with *Pst* and *Bgt,* respectively, were performed using tandem mass tags (TMT) technology. A total of 741 proteins were identified as differentially accumulated proteins (DAPs). Bioinformatics analyses indicated that some functional categories, including antioxidant activity, exhibited obvious differences between *Pst* and *Bgt* infections. Intriguingly, only 42 DAPs responded to both *Pst* and *Bgt* infections. Twelve DAPs were randomly selected for RT-qPCR analysis, and the mRNA expression levels of eleven were consistent with their protein expression. Furthermore, gene silencing using the virus-induced gene silencing system indicated that glutathione S-transferase (TaGSTU6) has an important role in resistance to *Bgt* but not to *Pst*. TaGSTU6 was shown to interact with the cystathionine beta-synthase (CBS) domain-containing protein (TaCBSX3). Knockdown of *TaCBSX3* expression only reduced wheat resistance to *B*g*t* infection. Overexpression of *TaGSTU6* and *TaCBSX3* in Arabidopsis (*Arabidopsis thaliana*) promoted plant resistance to *Pseudomonas syringae* pv. *tomato* DC3000 (*Pst* DC3000). Our results indicated that the TaGSTU6 interacting with TaCBSX3 only confers wheat resistance to *Bgt*, suggesting that wheat has different response mechanisms to *Pst* and *Bgt* stress.

**One-sentence summary:** Proteomics revealed a difference in the wheat resistance response to *Pst* and *Bgt*, and the TaGSTU6/TaCBSX3 interaction plays an important role only in wheat resistance to *Bgt*.

## Introduction

Wheat is one of the main human foodstuffs and is planted extensively around the world. Wheat stripe rust and powdery mildew, caused by the biotrophic pathogens, *Puccinia striiformis* f. sp. *tritici* (*Pst*) and *Blumeria graminis* f. sp. *tritici* (*Bgt*), respectively, are two of the most important foliar diseases worldwide. In China, these two diseases cause enormous losses of wheat yield and quality (Wellings, 2011). Reasonable breeding and use of resistant wheat cultivars are the most economical, effective, and environment-friendly methods of controlling stripe rust and powdery mildew (Chen, 2005; Nelson et al., 2018). However, long-term survival pressure and environmental stimuli endow pathogens with superior adaptability (Turrà et al., 2014), which results in high variation in the *Pst* and *Bgt* (Lei et al., 2017). Resistance genes of wheat also become less effective as pathogens evolve, thus stripe rust and powdery mildew continue to threaten global wheat production and food security (Huang et al., 1997). Therefore, it is of great importance to study and explore new resistant genes to stripe rust and powdery mildew in wheat to devise long-term prevention and control of these diseases.

Transcriptome surveys of the wheat line N9134 inoculated with *Bgt* and *Pst* found that *Bgt* triggered a higher number of genes and pathways compared with *Pst* (Zhang et al., 2014). The Chinese winter wheat cultivar XM318 is highly resistant to all the current prevalent Chinese *Bgt* and *Pst* races and pathotypes. However, whether XM318 responds to these two obligate biotrophic fungi by similar resistance mechanisms is unknown. To date, tandem mass tags (TMT) technologies that have been developed to quantify proteins and drawn increasing attention (Qiao et al., 2021). Although TMT technology has been used in qualitative and quantitative protein researches in wheat-*Pst* and wheat-*Bgt* interactions (Yang et al., 2016; Li et al., 2017), no thorough proteomics studies have been carried to compare the interactions of wheat-*Pst* and wheat-*Bgt*. Therefore, in this study, proteomics analyses were performed to explore response difference of XM318 to *Pst* and *Bgt* interactions. Eventually, the glutathione S-transferase TaGSTU6 aroused our attention, because it was down-regulated in XM318-*Pst* and up-regulated in XM318-*Bgt*.

Glutathione S-transferase (GST) represents a group of ubiquitous proteins in plants that comprise various functional proteins. Wang et al. (2019) categorized plant GSTs into eight classes: tau (GSTU), phi (GSTF), lambda (GSTL), dehydroascorbate reductase (DHAR), theta (GSTT), γ-subunit of translation elongation factor (EF1G), zeta (GSTZ), and tetrachlorohydroquinone dehalogenase (TCHQD) (Wang et al., 2019). One of the most important functions of GSTs is their ability to inactivate toxic compounds. Soluble GSTs mediate degradation of reduced glutathione or its homologs that form complex with herbicides (Jablonkai and Hatzios, 1993). Furthermore, a lot of studies indicated that GSTs are involved in secondary metabolism (Mueller et al., 2000), growth and development (Gong et al., 2005), and biotic and abiotic stress response of plants (Pan et al., 2018).

In barley, GST has been proved to be interacted with the *Blumeria* effector BEC1054 which may compromise well-known key player of the defense and response to pathogen (Pennington et al., 2016). In addition, TaGSTU61 associates with the WRKY74 regulatory factor to mediate copper tolerance by affecting glutathione accumulation (Li et al., 2020). GST activity in emmer wheat leaves also contributes to decreased oxidative stress and increased plant resistance to herbicide agents (Karpenko et al., 2019). *TaGSTU6U1* and *TaGSTU6F6* are important for monocarpic senescence and drought stress (Secenji et al., 2009). However, to the best of our knowledge, the regulatory mechanisms whereby the *TaGSTU6* responds to fungal stresses, especially to *Pst* and *Bgt*, remain unclear. Analyzing the resistance mechanism of *TaGSTU6* will enable us to understand how wheat responds to different fungal infections.

In this study, based on quantitative proteomics analysis and DAP identification between wheat XM318-*Pst* and XM318-*Bgt* interactions. We have systematically analyzed the relationships between the resistance mechanisms of wheat to *Pst* and *Bgt*, and have identified differences in protein regulation in these interactions. Furthermore, we have identified *TaGSTU6*, which is regulated at the protein and mRNA level during *Pst* and *Bgt* infections. Knockdown of *TaGSTU6* expression reduced wheat resistance to *Bgt,* but had no obvious effect on *Pst* resistance. Further analyses showed that TaCBSX3, as a target of TaGSTU6, is a positive regulator of wheat resistance to *Bgt*. In addition, overexpression of *TaGSTU6* or *TaCBSX3* in Arabidopsis promoted plant resistance to *Pst* DC3000. The results provide a new foundation to broaden our understanding of the molecular mechanism of the wheat**-***Pst* and wheat**-***Bgt* interactions at protein level and to provide a theoretical basis for resistance breeding and sustainable control of stripe rust and powdery mildew.

## Results

### Important Time Points for *Pst* and *Bgt* Resistance in XM318 are 24 and 48 Hours Post Inoculation (hpi)

In a study of histologically observation, we observed that the wheat cultivar XM318 was highly resistant to *Pst*, and immune to *Bgt*. Full rust disease symptoms were developed at 16 days post inoculation (dpi) in the susceptible control MX169, but XM318 displayed only slight necrotic flecks (**Fig. 1A**). For *Bgt* infections, the leaves of XM318 failed to develop obvious signs of infection and exhibited remarkable powdery mildew resistance at 12 dpi compared to the abundant surface mycelia in the susceptible control JS16 (**Fig. 1A**).

**Figure 1.**
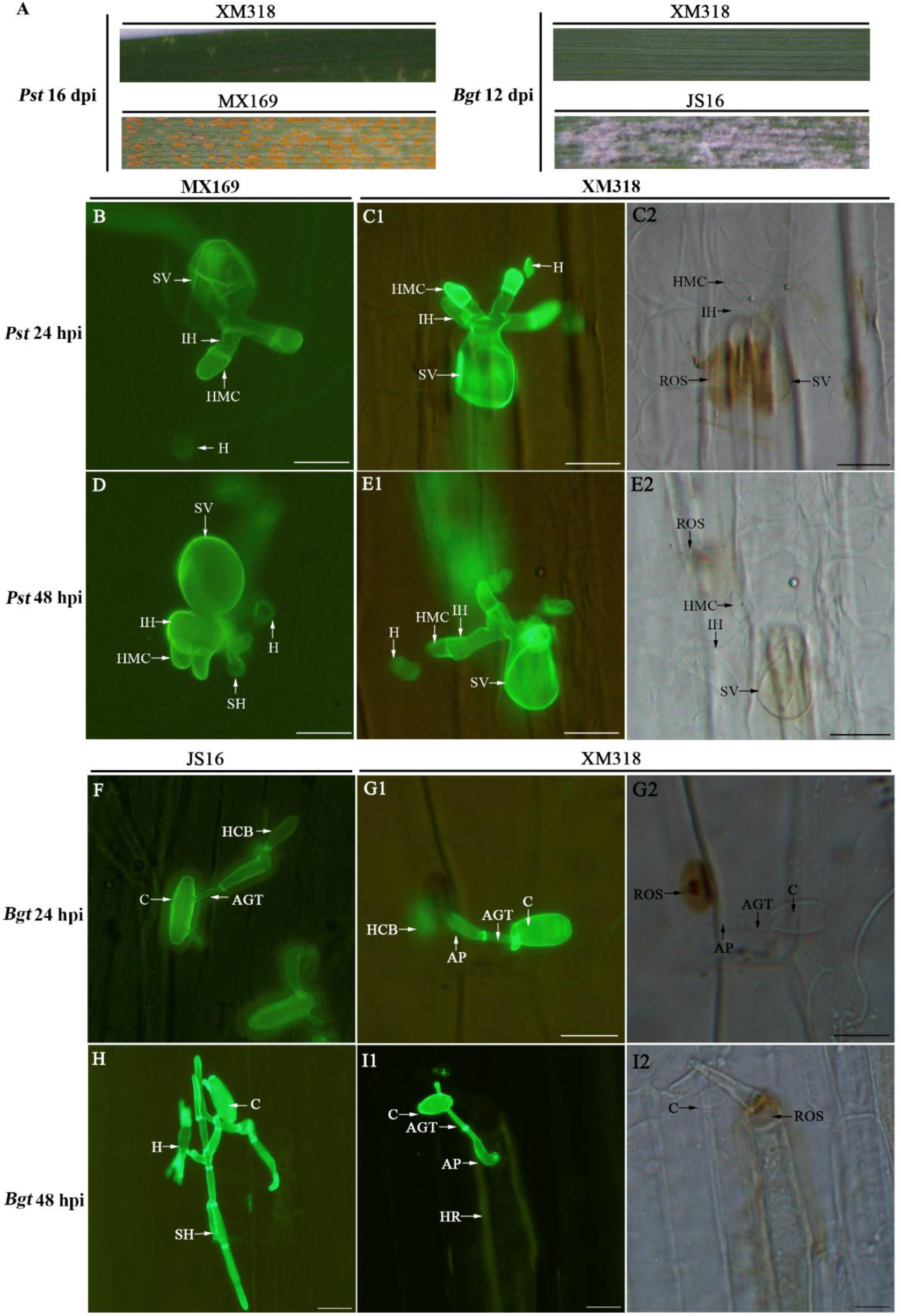
Epifluorescence observations of fungal growth in the wheat cultivars XM318 and MX169 infected with *Pst*, and XM318 and JS16 infected with *Bgt*. Phenotypes of XM318 and MX169 wheat cultivars infected with Pst observed at 16 dpi, and phenotypes of XM318 and JS16 wheat cultivars infected with Bgt observed at 12 dpi (A). Leaves sampled at 24 and 48 hpi and examined by epifluorescence microscopy after staining with WGA conjugated to Alexa-488. MX169 (B) and XM318 (C1, C2) inoculated with *Pst* at 24 hpi. MX169 (D) and XM318 (E1, E2) inoculated with *Pst* at 48 hpi. JS16 (F) and XM318 (G1, G2) inoculated with *Bgt* at 24 hpi; JS16 (H) and XM318 (I1, I2) inoculated with *Bgt* at 48 hpi. Scale bars = 20 µm. C, conidia; AP, appressorium; AGT, appressorium germ tube; PGT, primary germ tube; HR, hypersensitive reaction; SV, substomatal vesicle; HMC, haustorial mother cell; IH, infection hypha; NHC, necrotic host cell; H, haustoria; HCB, haustoria central body; ROS, reactive oxygen species.

To determine the key time points of immune response of XM318 to the two obligate biotrophic parasitic fungi, the infection processes of *Pst* and *Bgt* were investigated histologically at 12, 24, 36, 48, 60, 72, 84, and 96 hours post inoculation (hpi). Our results showed that the hyphal length of *Pst* was significantly (P < 0.05) shorter in XM318 than in the susceptible control MX169 at 24 hpi, indicating that XM318 began to set up resistance at 24 hpi (**Fig. 1, B and C1; Supplemental Table S1-1).** At 48 hpi, the hypha length and area of infection of *Pst* were significantly reduced in XM318 in comparison with in MX169. Moreover, infected hyphae (IH) of the control inflated obviously and secondary hyphae (SH) was observed in MX169, but not in XM318 (**Fig. 1, D and E1; Supplemental Table S1-1**), suggesting that SH formation was restricted in XM318. Furthermore, the numbers of haustorial mother cells (HMC) and hyphae branches increased more slowly after 48 hpi in the XM318 plant cells than in MX169 (**Fig. S1; Supplemental Table S1-1**).

*Bgt* infection also resulted in haustorial center bodies (HCB) at 24 hpi in both XM318 and the susceptible control JS16 (**Fig. 1, F and G1**), but colony formation was inhibited in XM318 leaves (**Supplemental Table S1-2**). At 48 hpi in the JS16 control cultivar, the formation of mature haustoria, development and growth of SH, and fungal growth was significantly greater than that in the XM318 resistant leaves (**Fig. 1, H and I1; Supplemental Table S1-2**). Interestingly, in contrast to the susceptible control, the hyphae almost stopped growing, and the colonies began to abort as hypersensitive reactions became evident after 24 hpi in XM318 (**Fig. 1, I1; Fig. S2; Supplemental Table S1-2**). In total, these results suggest that 24 hpi is an important time point that represents the early stages of the XM318-*Pst* and XM318-*Bgt* interactions. Also, 48 hpi is an important time point, because secondary hyphae was observed in both JS16-*Bgt* and MX169-*Pst* susceptible combination, but was restricted in the XM318-*Pst* and XM318-*Bgt* combination. Thus, the onset of resistance in both XM318-*Pst* and XM318-*Bgt* appears to occur between 24 and 48 hpi during which secondary hyphae begin to appear in the primary infection foci. Therefore, these times merited attention in the subsequent experiments.

### H_2_O_2_ Accumulation in XM318-*Bgt* is Faster than in XM318-*Pst*

To elucidate the immune response kinetics of XM318 to *Pst* or *Bgt* infection, hydrogen peroxide (H_2_O_2_) accumulation and the percentage infection sites with necrosis were measured. H_2_O_2_ accumulation could be detected microscopically in the leaves infected with *Pst* or *Bgt* at 24 hpi (**Fig. 1, C2 and G2**), and it reached a peak at 48 to 72 hpi in both interactions and began to decline as the percentage of necrotic cells increased later in infection, but development of H_2_O_2_ in XM318-*Bgt* was more rapid than in XM318-*Pst* at 36 hpi (**Supplemental Table S2**). In addition, hypersensitive necrosis of mesophyll cells were observed at 36 hpi in leaves infected with *Pst* or *Bgt* (**Supplemental Fig. S1F2, Fig. S2F2 and Supplemental Table S2**). And the rate of necrosis site development in XM318-*Bgt* was higher than that of XM318-*Pst* (**Supplemental Table S2**). These results indicate that the response kinetic of XM318 to *Bgt* infection is different from that kinetics to *Pst* infection.

**Figure 2.**
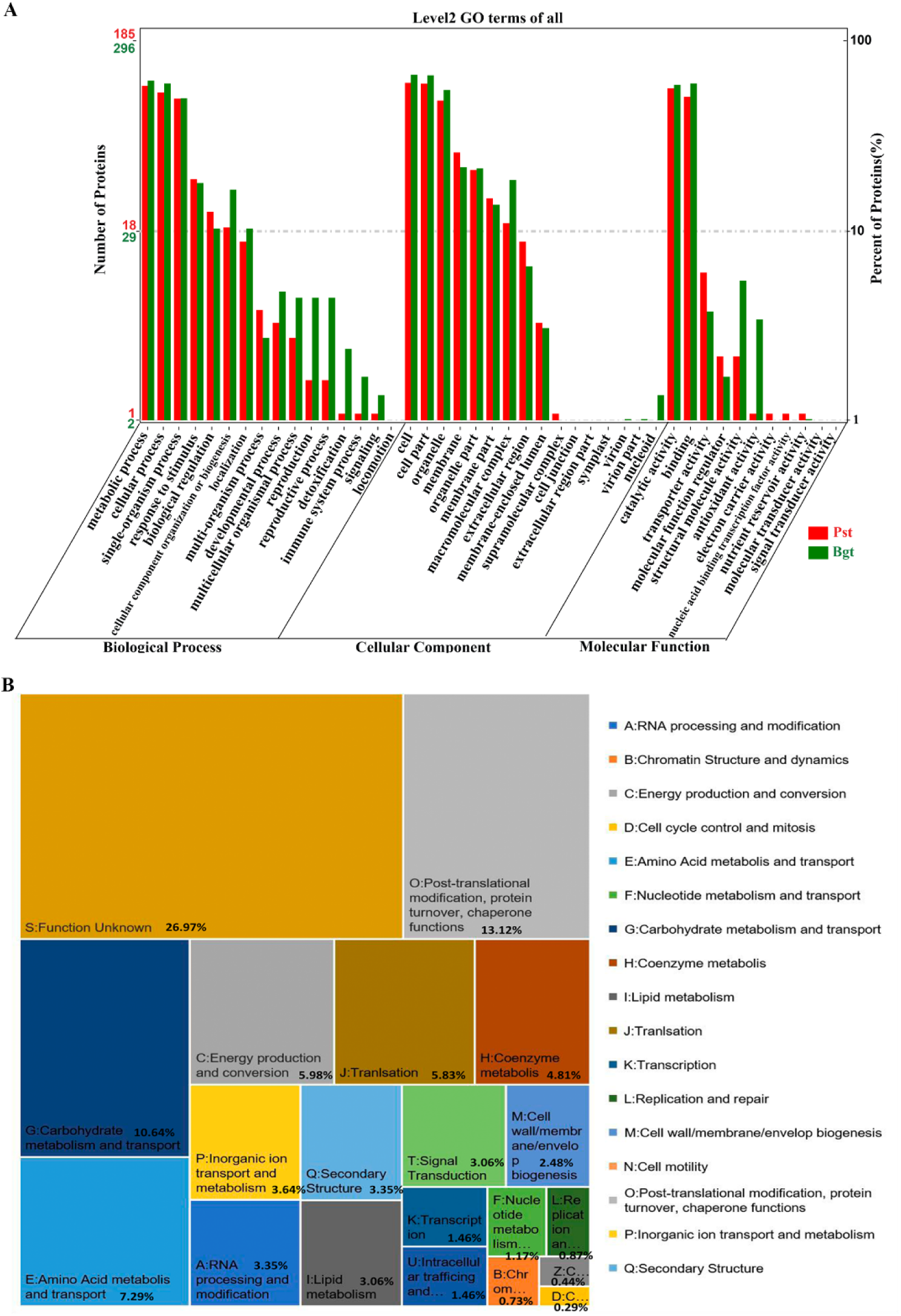
Functional analysis of DAPs. A, Multi-group data of GO enrichment analyses of HRPs during *Pst* and *Bgt* infections of wheat. *Pst* and *Bgt* indicate DAPs identified from wheat after *Pst* and *Bgt* infections. The numbers on the left Y axis represent the number of DAPs annotated in different functional categories. B, COG function classification of HRPs identified by TMT analyses of wheat infected with *Pst* and *Bgt*. A total of 501 proteins were annotated and assigned to 17 categories by COG analysis.

### Overview of Tandem Mass Tag (TMT) Data

To characterize the response difference between XM318 to *Pst* and XM318 to *Bgt* at protein level, proteomics analysis were performed on wheat leaves infected with *Pst* or *Bgt* at 24 hpi and 48 hpi, respectively. A total of 834,046 MS^2^ mass spectra were obtained (**Supplemental Table S3**). MS^2^ mass spectra were analyzed using the MASCOT software to obtain each MS^2^ mass spectrum score, which showed that the distribution of the MS^2^ mascot score is conformed to normal distribution (**Supplemental Fig. S3A**). The raw data was filtered and searched using the MASCOT engine and Proteome Discoverer, and the 171,645 total mass spectra were matched the number of peptides. Finally, a total of 36,854 matched peptides, 22,159 matched unique peptides, and 7,311 proteins were identified. The qualitative proteins were 7,311, and the quantified proteins were 5,485 (**Supplemental Table S3**). In addition, about 90% of the identified proteins had a molecular weight distribution between 10-110 kDa, which followed the normal distribution (**Supplemental Fig. S3B**). A total of 741 proteins were significantly different between treatments (*Pst* or *Bgt* infected at 24 hpi or/and 48 hpi) and identified as differentially accumulated proteins (DAPs). Moreover, all identified proteins and DAPs were investigated by heat map analysis, which provides a clear visual perception for a large amount of complex protein data (**Supplemental Fig. S4**).

### Bioinformatics Analysis of DAPs

All DAPs were derived from two categories: horizontal regulated proteins (HRPs: DAPs from comparisons of proteomic data from leaves infected with the same pathogen but collected at different time points) and vertical regulated proteins (VRPs: DAPs of comparative proteomic data from different pathogen-infected leaves at the same time points) (**Supplemental Fig. S5A**). Among 516 HRPs, 213 proteins (**Supplemental file S2**) were from *Pst* infections (0h vs *Pst*24h, 0h vs *Pst*48h), and 345 (**Supplemental file S3**) from *Bgt* infections (0h vs *Bgt*24h, 0h vs *Bgt*48h) (**Supplemental Fig. S5B**). There were 544 VRPs (*Pst*24h vs *Bgt*24h; *Pst*48h vs *Bgt*48h) (**Supplemental file S4**). Intriguingly, only 42 proteins (**Supplemental Table S4; Supplemental file S5**) were regulated in both *Pst* and *Bgt* infections, which led to further investigation. In *Pst*-induced DAPs, there were more up-regulated HRPs than down-regulated HRPs, contrary to *Bgt*-induced DAPs (**Supplemental Fig. S5C**). These findings suggest that XM318 responds to *Bgt* and *Pst* by implementing different proteins.

Gene Ontology (GO) and OmicShare Gene Ontology (OSGO) enrichment analyses showed that the HRPs were mainly categorized into cellular components and basal biological processes (**Fig. 2A, Supplemental Fig. S6**). OSGO analyses displayed that the number of DAPs belong to antioxidant activity and the immune system process were significantly higher in the XM318-*Bgt* than in the XM318-*Pst* interaction, consistent with the observation that the accumulation of H_2_O_2_ was more rapid during XM318-*Bgt* than XM318-*Pst*, and XM318 is immune to *Bgt* but high resistant to *Pst*. Although the number of DAPs belong to response to stimulus are similar between XM318-*Pst* and XM318-*Bgt* interactions, protein kinases and hormonal response proteins included in response to stimulus categories were substantially different between XM318-*Pst* and XM318-*Bgt* at both 24 hpi and 48 hpi (**Supplemental Table S5**). For example, the kinases MAPK5 and CDPK2 were up-regulated in XM318-*Bgt* but not in XM318-*Pst* interaction, and abscisic acid (ABA) and salicylic acid (SA) related proteins were more likely to be induced by *Pst*, and SA-, jasmonic acid (JA), and ABA-related proteins were mainly expressed in response to *Bgt*. The results indicated XM318 responded differentially to *Pst* and *Bgt* infections.

The Clusters of Orthologous Groups (COG) database was used to classify all DAPs (VRPs and HRPs). The main functional categories involved in post-translational modification (O, 90, 13.12%), carbohydrate metabolism and transport (G, 73, 10.64%), and other functional classification (**Fig. 2B; Supplemental file S6**). In summary, the COG analysis showed that the responses of XM318 to pathogen challenges comprise complex regulatory mechanisms.

To better understand the metabolism pathways involved in XM318-*Pst* and XM318-*Bgt*, plant visualization pathways and Kyoto Encyclopedia of Genes and Genomes (KEGG) pathway enrichment analyses were performed. KEGG analysis showed that lipid metabolism and plant-pathogen interactions were activated in XM318 upon *Pst* infection, suggesting their vital roles during XM318-*Pst* interaction. In the XM318-*Bgt* interaction, DAPs were significantly enriched in plant-pathogen interactions and MAPK signaling pathway, indicating their important roles (**Supplemental Fig. S8**). These data show a correlation in the XM318 response to the *Pst* and *Bgt* pathogens, suggesting that differences exist in the main metabolic pathways.

of 12 randomly selected HRPs.

### Gene Expression Analysis of Selected DAP Genes by RT-qPCR

RT-qPCR analyses were performed to further verify the gene expression profiles with *Pst* or *Bgt* infections. We selected 12 DAPs based on significant differential expression induced by *Pst* and *Bgt* together or alone. Among these DAPs, six proteins were regulated in both *Pst* and *Bgt* infections and the other six proteins accumulated differently in *Pst* or *Bgt* infections (**Supplemental Table S6**). The transcriptional expression levels of 11 genes were consistent with their protein levels from proteomic data for at least one sampling time point (**Supplemental Fig. S9**). However, the expression patterns of TaGSTU6 (UniProt database, A0A1D5RWE5) were consistent between the mRNA and protein levels in XM318-*Bgt* but inconsistent in XM318-*Pst*. RT-qPCR verified opposite gene expression patterns in *Pst* and *Bgt* infections, such as Caleosin and MODIFIRE OF SNC1 11 (**Supplemental Fig. S9, E and F**), suggesting that differences exist in the wheat responses to *Pst* and *Bgt* infections. In conclusion, the proteomics analysis provided more accurate and simpler data.

### Identification of TaGSTU6

Intriguingly, only 42 proteins (**Supplemental Table S4; Supplemental file S5**) were regulated in both *Pst* and *Bgt* infections, among which four were down-regulated in *Pst* infection but up-regulated in *Bgt*-infection. TaGSTU6 was selected for further study. RT-qPCR was performed to elucidate the transcript levels of *TaGSTU6* during wheat-*Pst* and wheat-*Bgt* interaction. The results revealed that the *TaGSTU6* transcripts were up-regulated during *Bgt* and *Pst* infections, and its transcript levels in compatible combinations were lower than that in the incompatible combinations (**Supplemental Fig. S10, A and B; Supplemental Table S7**). BLAST analysis on the hexaploid wheat genome databases revealed three *TaGSTU6* copies located on chromosomes 1A, 1B, and 1D, respectively. Phylogenetic analysis showed that TaGSTU6 is closely related to HvGSTU6 (*Hordeum vulgare*; 92.27%) and OsGSTU6 (*Oryza sativa*; 70.39%) (Supplemental Fig. S10C). Sequence analyses indicated that TaGSTU6 contains a typical GST C-terminal domain and an N-terminal domain that bearing a transmembrane helix (**Supplemental Fig. S10D**).

### Silencing of *TaGSTU6* Attenuates Wheat Resistance to *Bgt*

To evaluate the function of TaGSTU6 during wheat-*Pst* or wheat-*Bgt* interaction, the barley stripe mosaic virus (BSMV)-virus-induced gene silencing (VIGS) system was used to silence *TaGSTU6* in XM318. At 10 days post BSMV inoculation, BSMV:*TaPDS* plants exhibited a bleaching phenotype, and mild chlorotic mosaic symptoms were observed in almost all BSMV-inoculated plants (**Fig. 3A**), indicating that the BSMV-VIGS system was functional and that virus inoculations were successful and stable.

**Figure 3.**
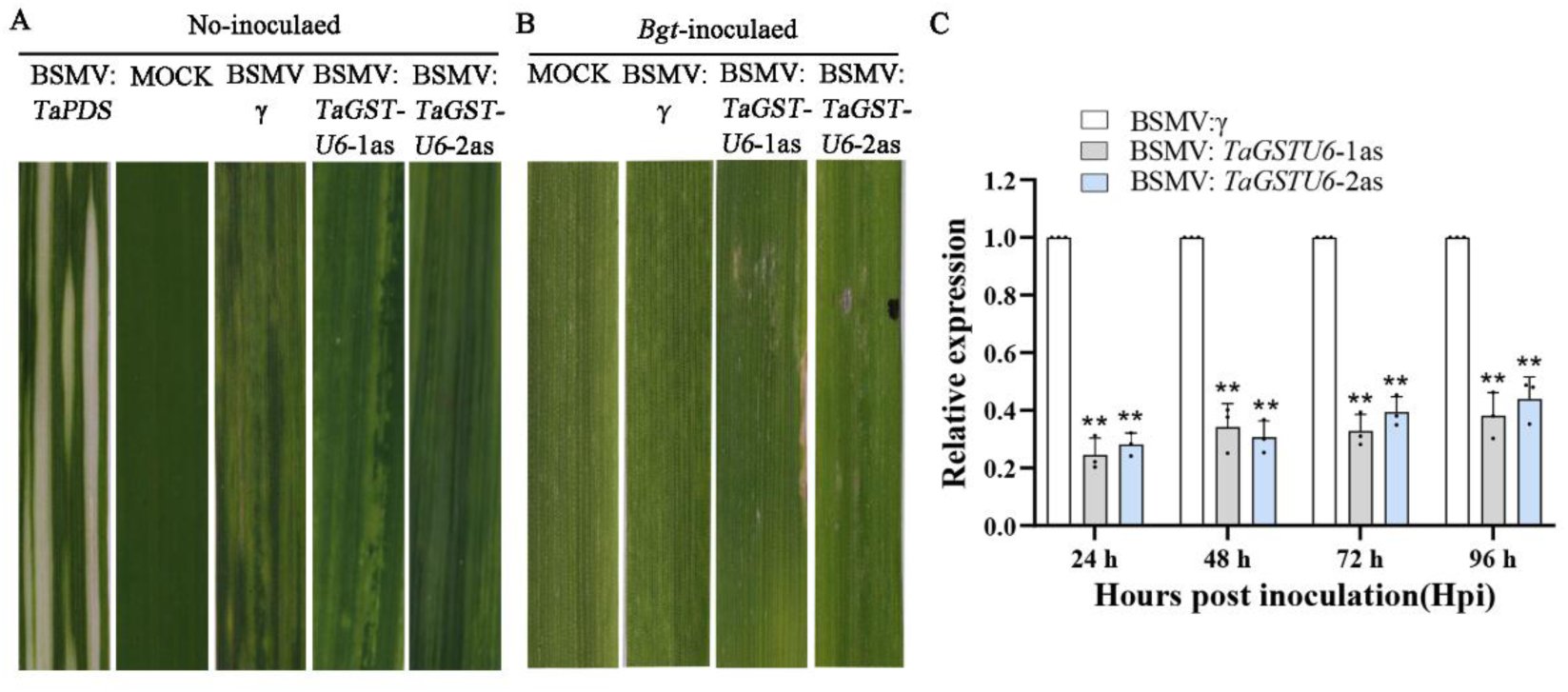
Functional assessment of *TaGSTU6* in the wheat-*Bgt* incompatible interaction by VIGS. A, Phenotypes of XM318 wheat leaves inoculated with BSMV:*TaPDS*, FES buffer (mock), BSMV:γ, and BSMV:*TaGSTU6* at 10 dpi. B, Phenotypes of the fourth emerging wheat leaves at 12 dpi after *Bgt* infection. Plants were pre-inoculated with FES buffer (mock), BSMV:γ, or BSMV:*TaGSTU6*. C, Relative transcript levels of *TaGSTU6* in silenced plants inoculated with *Bgt* at 0, 24, 48, and 96 hpi. Wheat leaves inoculated with BSMV:γ and sampled after inoculation with *Bgt* were used as controls. Data are the means ± SE of three independent samples. Asterisks indicate significant differences at the same time points using Student’s t test (**P < 0.01).

The XM318 inoculated with BSMV:*TaGSTU6* (*TaGSTU6*-1as and *TaGSTU6*-2as) were greatly reduced in their resistance to *Bgt* in comparison with the control (XM318 inoculated with BSMV:γ) (**Fig. 3B**), while inoculation with BSMV:*TaGSTU6* had no obvious effects on the resistance of XM318 to *Pst* (**S**upplemental Fig. S11 and Fig. S12). When compared transcript levels between the BSMV:*TaGSTU6* inoculated leave with the control leaves, the *TaGSTU6* transcripts were reduced by 62–75% for *TaGSTU6*-1as and by 67–76% for *TaGSTU6*-2as (**Fig. 3C**), indicating that the reduced resistance of XM318 to *Bgt* could be due to the silence of *TaGSTU6.* WGA staining and confocal microscopy observation revealed that although no significant hyphal growth differences were observed between the *TaGSTU6*-silenced and control plants at 24 hpi (Fig. 4, A and B), *Bgt* progressed much faster in the *TaGSTU6*-silenced plants than in the control at 48 hpi, 96 hpi, and 12 dpi (**Fig. 4, C-H**). The number of hyphal branches/infection, HMC, hyphal length, and total infection areas were significantly higher in the silenced leaves than in control leaves at 48 and 96 hpi (**Fig. 4, I-L**). These results indicated that TaGSTU6 played an important role in XM318 resistance to *Bgt* after 48 hpi. To further verify its function in wheat resistance to *Bgt*, *TaGSTU6* was also silenced in the susceptible wheat-*Bgt* compatible interaction. Aggravated disease phenotypes and larger colony morphologies were observed in *TaGSTU6*-knockdown plants at 12 dpi (**Supplemental Fig. S13 and Fig. S14**). In conclusion, *TaGSTU6* plays a positive role in wheat resistance to *Bgt*, but has no obvious disease-resistance functions during wheat-*Pst* infection.

**Figure 4.**
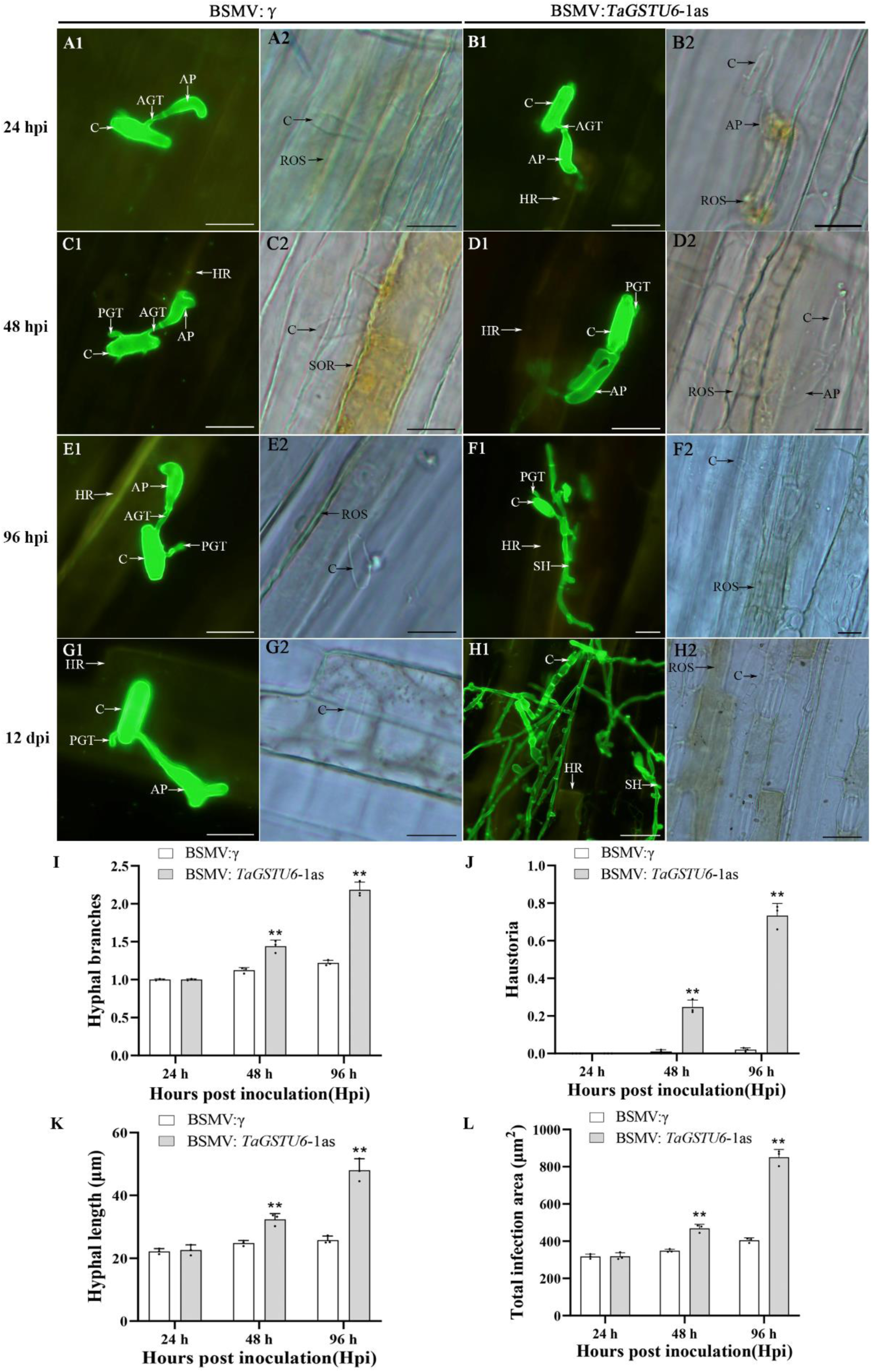
Epifluorescence observations of fungal growth in the XM318 wheat cultivar inoculated with BSMV and infected with *Bgt* (incompatible combination). A-G, Fungal growth examined under an epifluorescence microscope after staining with WGA conjugated to Alexa-488. Inoculation with BSMV:γ or BSMV:*TaGSTU6* was carried out and followed after 12 dpi by *Bgt* infection. Images were then captured at 24 hpi (A, a and B, b), 48 hpi (C, c and D, d), 96 hpi (E, e and F, f), and 12 dpi (G, g and H, h). The same letter indicates the photo was taken at the same infection site. I-L, The number of hyphal branches (I), haustoria (J), hyphal lengths (K), and total infection areas (L) were measured in wheat leaves at 24, 48, 96 hpi, and 12 dpi after inoculation with *Btg*. All results were obtained from 50 infection sites. Data are the means ± SE of three independent samples and differences were assessed using Student’s *t* tests. \**P* < 0.05; \*\**P* < 0.01. (A-G) scale bars = 20 µm; (H, h) scale bars = 50 µm.

### TaGSTU6 Interacts with the CBS Domain-containing Protein TaCBSX3

To identify the interacting proteins of TaGSTU6 during XM318-*Bgt* interaction, the cDNA of *TaGSTU6* was used as the bait to screen a yeast two-hybrid (Y2H) library constructed with RNA isolated from *Bgt*-infected XM318 leaves. A total of 11 clones representing eight XM318 proteins were screened out as potential TaGSTU6-interacting proteins (**Supplemental Table S8**), among which, the CBS domain-containing protein TaCBSX3 (XM_020317884.1) displayed the strongest interaction and was selected for subsequent studies. Sequence analyses revealed that TaCBSX3 contains two conservative CBS domains with three copies in the wheat genome that are located on chromosomes 4A, 4B, and 4D (the sequence identity of the three copies of TaCBSX3 gene is 97%, 99%, and 100% when compared to the *TaCBSX3* in chromosome 4D). Furthermore, when the TaGSTU6-GFP and TaCBSX3-mCherry constructs were coinfiltrated into *N. benthamiana* leaves, strong green and red fluorescence were co-localized in the plasma membrane and cytoplasm (**Fig. 5A**), suggesting the possible interaction of TaGSTU6-GFP and TaCBSX3-mCherry.

**Figure 5.**
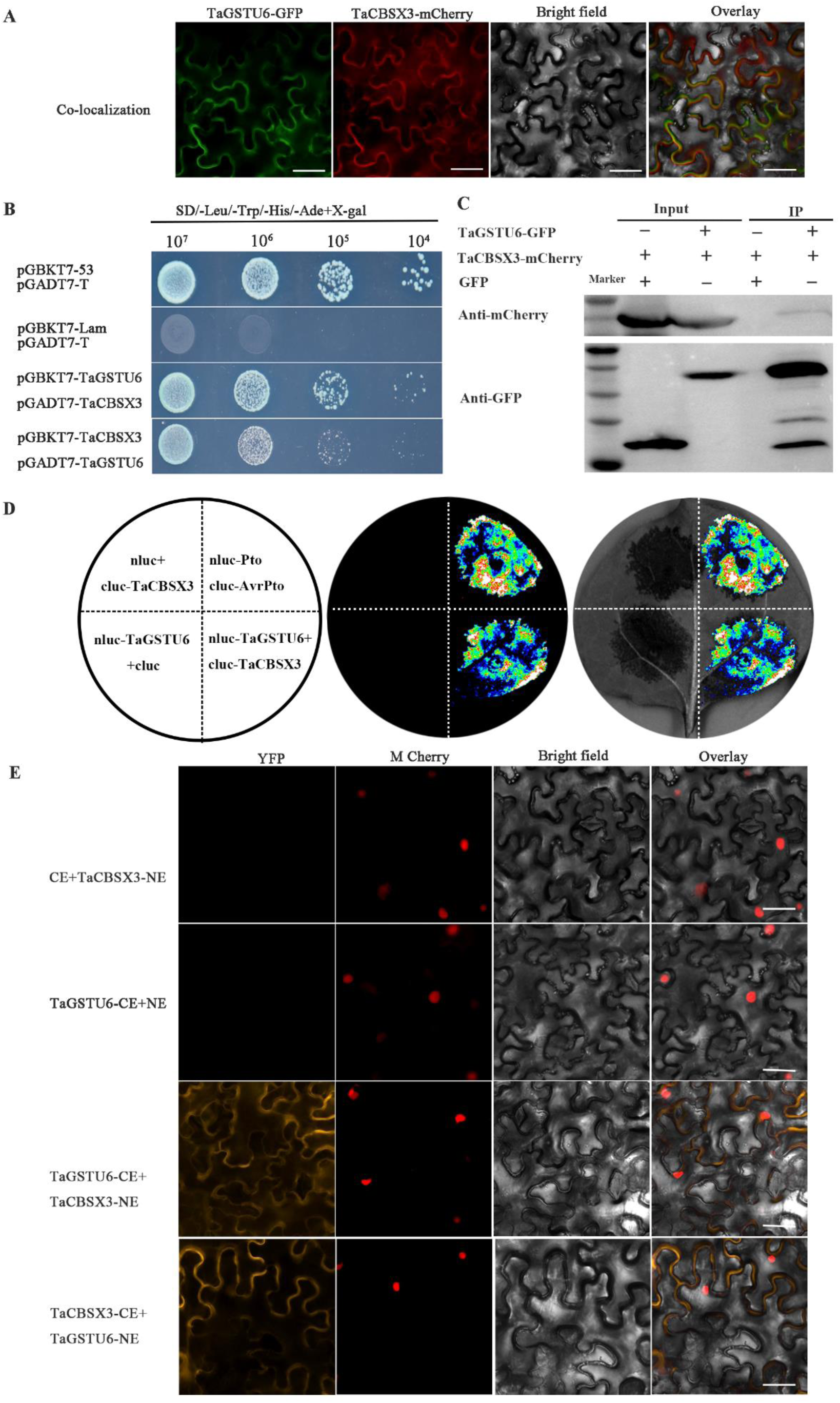
TaGSTU6 interacts with TaCBSX3 *in vivo*. A, Y2H analysis of TaGSTU6 interactions with TaCBSX3. Y2H Gold yeast cells transformed with the bait and prey vectors were assayed for growth on the selective medium SD/-Leu-Trp-His-Ade/X-α-gal/AbA. Yeast co-expressing TaGSTU6 and TaCBSX3 grew on the selective medium and yielded X-α-gal activity. Yeast cells containing pGADT7-T and pGBKT7-53 or pGBKT7-Lam vectors were used as positive and negative controls. B, Co-IP assay was performed to detect TaGSTU6-TaCBSX3 binding in plant cells. Western blots of total proteins from *N. benthamiana* leaves transiently expressing constructs carrying TaGSTU6-GFP and TaCBSX3-mCherry. Proteins eluted from GFP-Trap agarose beads were detected with anti-GFP or anti-mCherry antibodies. C, Confirmation of interactions between TaGSTU6 and TaCBSX3 by LCI assays in *N. benthamiana. N. benthamiana* leaves were infiltrated with *Agrobacterium* strains containing the indicated constructs. Signals were collected 48 h after infiltration. D, TaGSTU6 and TaCBSX3 interactions *in planta* detected by fluorescence complementation assays. *N. benthamiana* leaves were infiltrated with *Agrobacterium* strains containing the following construct pairs, CE+TaCBSX3-NE, TaGSTU6-CE+NE, TaGSTU6-CE+TaCBSX3-NE, and TaCBSX3-CE+TaGSTU6-NE. The infiltrated leaves were observed via fluorescence microscopy. The AtH2B histone protein was used as a nuclear localization marker gene. Scale bars = 20 µm.

We further confirmed the TaGSTU6-TaCBSX3 interaction with Y2H, co-immunoprecipitation (Co-IP), firefly luciferase complementation imaging (LCI), and bimolecular fluorescence complementation (BiFC) assays. Only yeast cells expressing both TaGSTU6 and TaCBSX3 grew on the selective medium SD/-Leu-Trp-His-Ade/X-α-gal/AbA and exhibited beta-galactosidase activity (**Fig. 5B; Supplemental Fig. S15**). In Co-IP assays with *N. benthamiana* leaves infiltrated with the TaGSTU6-GFP and TaCBSX3-mCherry fusion constructs, bands representing TaGSTU6-GFP and TaCBSX3-mCherry were present in the western blots of the total proteins and the proteins eluted from anti-GFP beads (**Fig. 5C**), indicating TaGSTU6 interacting with TaCBSX3 *in vivo*. In LCI assays, strong luciferase activity was observed with the nLuc-TaGSTU6 and cLuc-TaCBSX3 combinations, providing further evidence of the TaGSTU6-TaCBSX3 interactions in *N. benthamiana* (**Fig. 5D**). In addition, BiFC assays were performed in *N. benthamiana* leaves to detect TaGSTU6-TaCBSX3 interactions. Strong yellow fluorescence signals were observed in cells co-transformed with nEYFP-TaGSTU6 and cEYFP-TaCBSX3 (**Fig. 5E**). In summary, the results of Y2H, Co-IP, LCI, and BiFC provided strong evidence for TaGSTU6-TaCBSX3 interaction.

### TaCBSX3 Contributes to Wheat Resistance to *Bgt*

To further determine whether TaCBSX3 is involved in wheat resistance to *Bgt* and *Pst*, we first analyzed the expression patterns of *TaCBSX3* in the incompatible and compatible combinations between *Bgt/Pst* and wheat by RT-qPCR. The transcripts of *TaCBSX3* were upregulated in the incompatible *Bgt-*XM318 interaction, but down-regulated in the compatible combination (**Supplemental Fig. S16A; Supplemental Table S7**), suggesting that *TaCBSX3* may play a role in the resistance of XM318 to *Bgt*. Similar expression patterns but to a lower extent of it were observed in wheat-*Pst* interactions (**Supplemental Fig. S16B**), indicating that *TaCBSX3* may not be involved in the resistance of XM318 to *Pst*.

Subsequently, the expression of *TaCBSX3* was knocked down by BSMV-VIGS in the incompatible XM318-*Bgt* combination, and 10 days after virus inoculation, the plants displayed chlorotic mosaic symptoms (**Fig. 6A**). After inoculation with *Bgt*, sporadic single spore foci appeared on leaves of the *TaCBSX3* silenced plants at 12 dpi, whereas conidium failed to appear on the control and BSMV:γ unsilenced control plants (**Fig. 6B**). The silencing efficiency confirmed that endogenous *TaCBSX3* expression was reduced in the silenced plants by ∼50 to 86% compared with the control plants (**Fig. 6C**). However, after inoculation with *Pst*, BSMV:*TaCBSX3* inoculated XM318 failed to exhibit obvious phenotypic differences compared with BSMV:γ inoculated control plants (**Supplemental Fig. S17 and Fig. S18**). Therefore, both the expression profile of *TaCBSX3* and the results of BSMV-VIGS experiment suggested that *TaCBSX3* played an important role in wheat resistance to *Bgt* but not to *Pst*.

**Figure 6.**
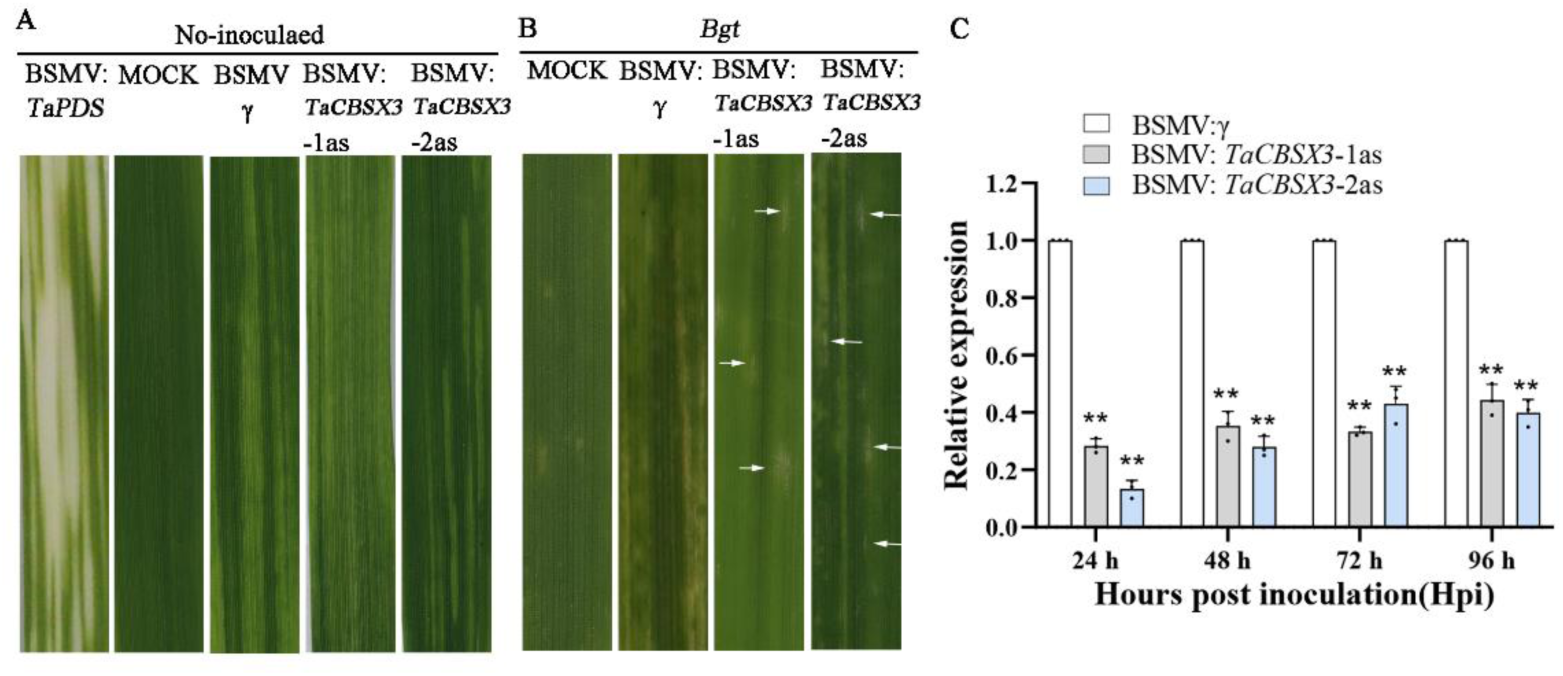
BSMV-induced gene silencing of *TaCBSX3* reduces wheat resistance against *Bgt* (incompatible combination). A, Phenotypes of XM318 wheat leaves inoculated with BSMV:*TaPDS*, FES buffer (mock), BSMV:γ, and BSMV:*TaCBSX3* at 10 dpi. B, *Bgt* infection phenotypes on the fourth leaves at 12 dpi. C, Relative transcript levels of *TaCBSX3* in silenced plants inoculated with *Bgt* at 0, 24, 48, and 96 hpi. Control wheat leaves were inoculated with BSMV:γ and sampled after inoculation with *Bgt*. Data are the means ± SE of three independent samples. Asterisks indicate significant differences at the same time points using Student’s t test (**P < 0.01). The arrow indicate the *Bgt* conidium.

Although no substantial histological differences between the *TaCBSX3*-knockdown and control plants were observed microscopically at 24 hpi (**Fig. 7, A-C**), the resistance of the *TaCBSX3*-silenced plants to *Bgt* was reduced at 48, 72, and 96 hpi, in comparison with that of the control plants (**Fig. 7, D-L**), indicating that *TaCBSX3* functions at 48 hpi. Statistical analyses also confirmed that the number of *Bgt* hyphal branches, hyphal length, and the total infected areas remarkably increased in the BSMV:*TaCBSX3*-inoculated leaves at 48, 72, and 96 hpi (**Fig. 7, M-O**). Furthermore, the necrotic areas of the *Bgt* infected leaves of the *TaCBSX3*-silenced plants were significantly smaller than the BSMV:γ-inoculated plants after 48 hpi (**Fig. 7P**), suggesting that *TaCBSX3* is important for the resistance of XM318 to *Bgt*. In order to verify the disease resistance effects of *TaCBSX3*, VIGS assays were performed to silence *TaCBSX3* in the wheat-*Bgt* compatible interaction, and the BSMV:*TaCBSX3*-inoculated leaves also appeared to be more susceptible to *Bgt* (**Supplemental Fig. S19 and Fig. S20**). Together, our results indicated that TaCBSX3 positively regulates wheat resistance to *Bgt*.

**Figure 7.**
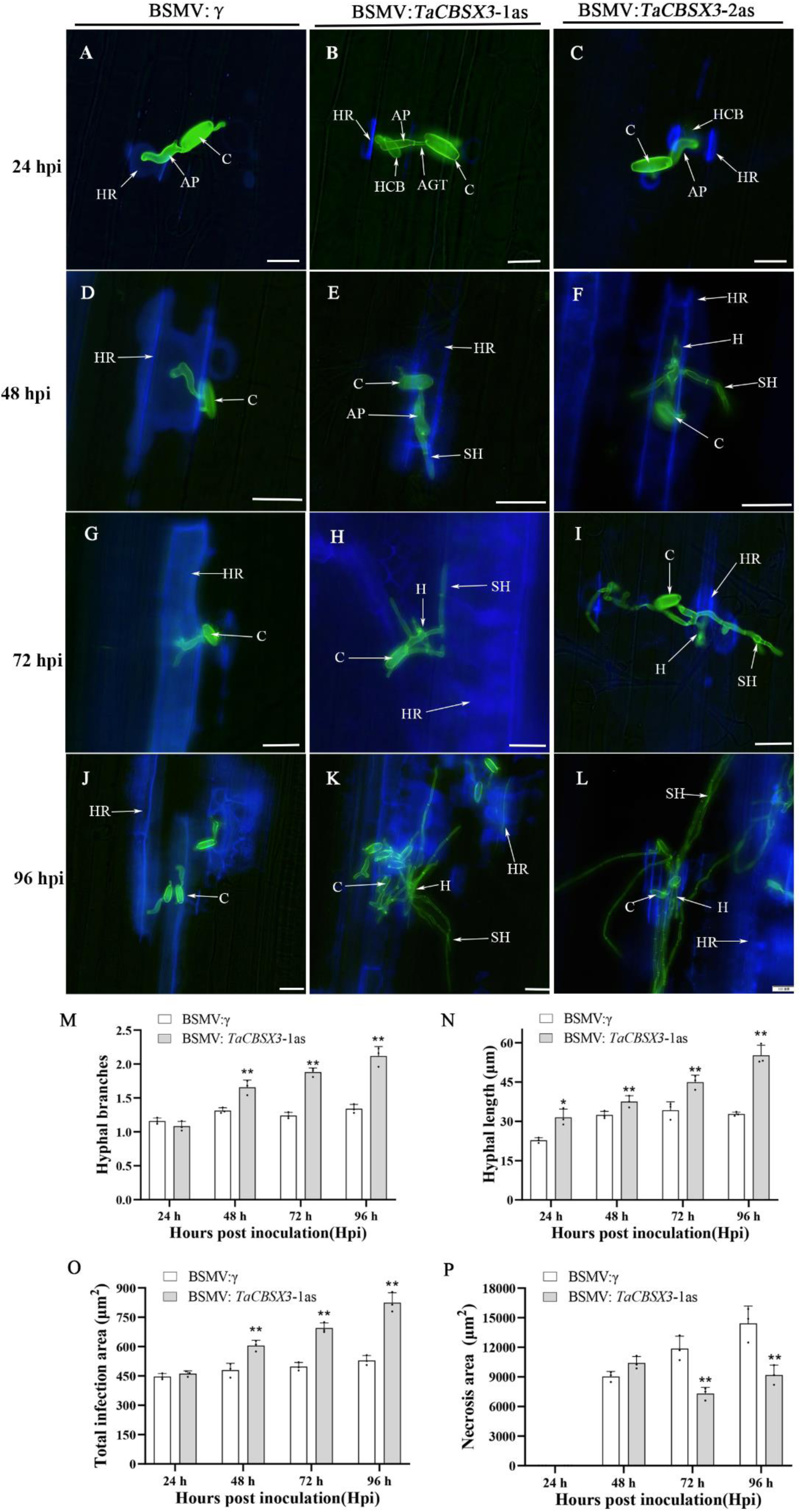
Histological observations of fungal growth in XM318 inoculated with BSMV after infection with incompatible *Bgt* combinations. A-L, Fungal structures on leaves were observed with an epifluorescence microscope after staining with WGA conjugated to Alexa-488. Inoculation with BSMV:γ or BSMV:TaCBSX3-1/2as and *Bgt* at 24 hpi (A-C), 48 hpi (D-F), 72 hpi (G-I), and 96 hpi (J-L). M-P, The numbers of hyphal branches (M), hyphal length (N), total infected areas (O) and necrotic areas (P) in BSMV-infected plants inoculated with *Bgt* were calculated with SPSS software. All results were obtained from 50 infection sites. Data are the means ± SE of three independent samples. Differences were assessed using Student’s t tests. \**P* < 0.05; \*\**P* < 0.01.

### Salicylic Acid (SA) Accumulation is Important for TaGSTU6 and TaCBSX3 Mediated Resistance to *Bgt* Infection

Plant hormones have been demonstrated to be critical to immune signaling processes (Vlot et al., 2009). And a bunch of hormone-related proteins in XM318 showed differential expression upon infection by *Pst* and *Bgt* in proteomic analysis. Therefore, we speculate that the endogenous hormone levels in XM318 varied between XM318-*Pst* and XM318-*Bgt*. To test this hypothesis, the endogenous levels of ABA, JA, SA, and ACC (1-aminocyclopropane-1-carboxylic acid: It is ethylene precursors or a plant hormone) in XM318 in response to *Pst* or *Bgt* were measured. The concentrations of ABA and SA in XM318 leaves were significantly up-regulated by *Pst* infection at 8 and 12 hpi compared with control (**Fig. 8A**). However, the SA was increased significantly in *Bgt*-infected leaves at 12 hpi, and ACC was increased significantly both 8 and 12 hpi (**Fig. 8B**). These results suggest that SA accumulation is important for wheat against both *Pst* and *Bgt* infections.

**Figure 8.**
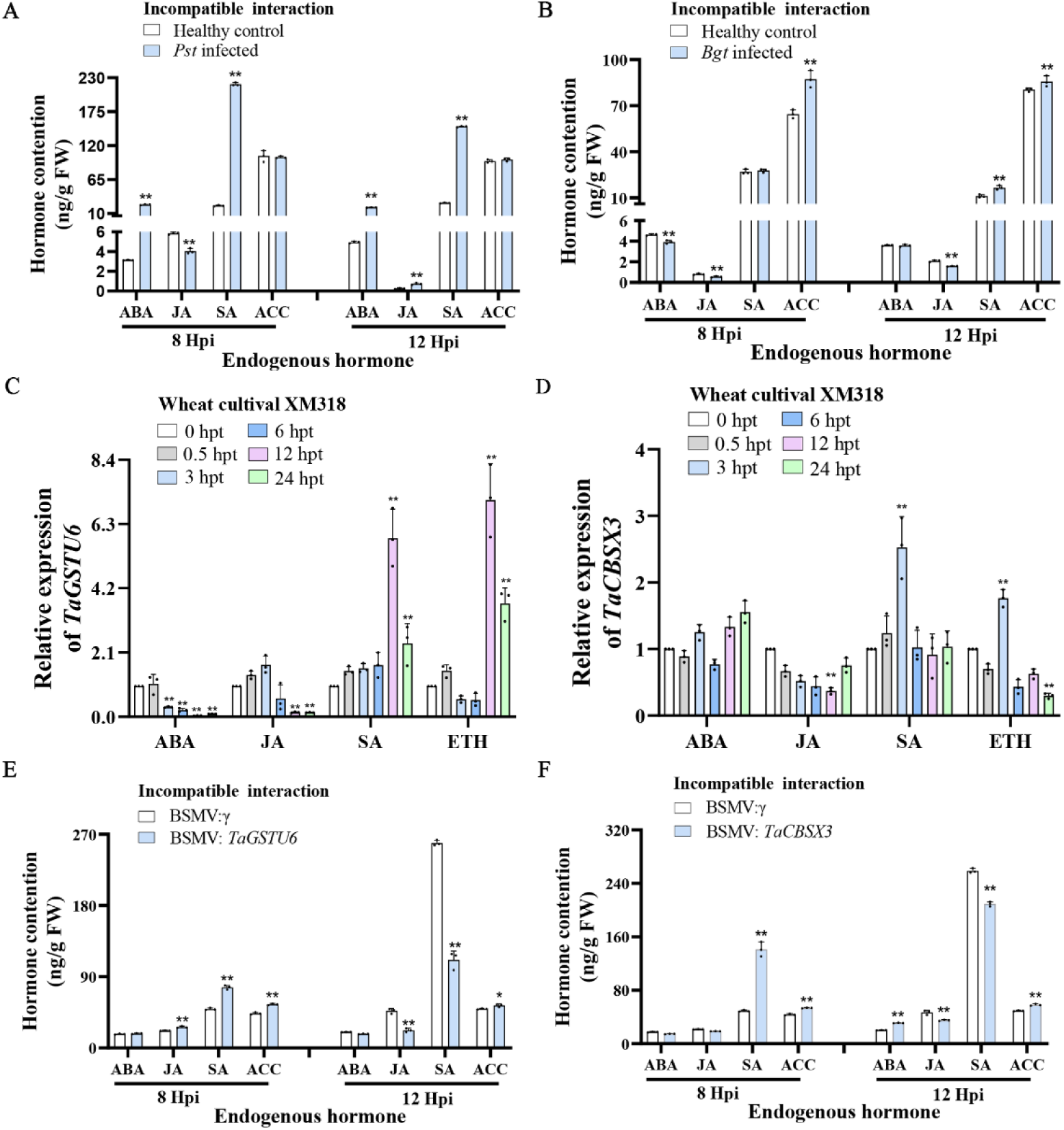
Expression profiles of *TaGSTU6* and *TaCBSX3* in response to hormones and the content of endogenous hormones in leaves in response to *Bgt* and *Pst* infections. A and B, RT-qPCR analyses of *TaGSTU6* and *TaCBSX3* expression after exogenous hormone treatments. ABA, abscisic acid; SA, salicylic acid; JA, jasmonate; ACC, 1-aminocyclopropane-1-carboxylic acid. Wheat seedlings treated with 0.1% (v/v) water were used as a mock control. Expression levels were normalized to *TaEF-1a*. C and D, Analyses of endogenous hormone content in wheat leaves inoculated with *Pst* or *Bgt*. ETH, ethylene. E and F, Hormone accumulations in non-silenced and *TaGSTU6*-*and TaCBSX3*-silenced leaves after inoculated with *Bgt*. Data are the means ± SE of three independent samples. Differences were assessed using Student’s t tests. \**P* < 0.05; \*\**P* < 0.01. FW, fresh weight.

To determine whether their expression were regulated by plant hormones, the transcription profiles of *TaGSTU6* and *TaCBSX3* in response to exogenous hormones were analyzed in resistant and susceptible cultivars by RT-qPCR, respectively. The transcript levels of *TaGSTU6* significantly increased in wheat leaves after SA or ETH treatments, and also had a positive response to JA and ABA treatment (**Fig. 8C; Supplemental Fig. S21A**). The expression levels of *TaCBSX3* also increased after SA and ETH treatment, and the abundance of the *TaCBSX3* transcripts decreased after JA treatment (**Fig. 8D; Supplemental Fig. S21B**). Therefore, the transcription of *TaGSTU6* and *TaCBSX3* is regulated by exogenous applied hormones.

To further explore the involvement of hormones in TaGSTU6 and TaCBSX3-related resistance to *Bgt* infection, the hormone accumulation in *TaGSTU6* and *TaCBSX3*-silenced and XM318 leaves upon *Bgt* infection were measured. Surprisingly, only SA accumulation was significantly reduced in the *TaGSTU6* or *TaCBSX3*-silenced leaves at 12 hpi, whereas the other hormones remained relatively stable or exhibited little impact after *TaGSTU6* or *TaCBSX3*-silencing (**Fig. 8, E and F**). Together, these results imply that SA has an important role for TaGSTU6 and TaCBSX3-mediated resistance to *Bgt* infection.

### Overexpression of *TaGSTU6* or *TaCBSX3* Enhances Arabidopsis Resistance to Pseudomonas syringae pv. tomato (Pst) DC3000

To verify the function of TaGSTU6, we overexpressed it in Arabidopsis challenged by *Pseudomonas syringae* pv. *tomato* (*Pst*) DC3000. The *TaGSTU6* overexpression constructs were introduced into Arabidopsis ecotype Columbia-0 (wild-type) and generated a set of transgenic lines (*TaGSTU6*-OE). BLASTp analysis showed that the AtGSTU17 (*Arabidopsis thaliana*, AY091102.1) is homologous of TaGSTU6. Also, the homozygous *atgstu1* mutants were obtained by screening subcultures from *atgstu*-F1 (SK88) treatments. Then, *TaGSTU6*-OE lines, homozygous *gstu* mutant lines, and wild-type (WT) Col-0 plants were challenged with *Pst* DC3000. Obvious chlorotic symptoms on the leaves of two *atgstu1* lines (*atgstu1*-9 and *atgstu1*-13) were observed at 2 dpi, indicating that *atgstu1* lines are more susceptible to *Pst* DC3000 than WT plants, indicating AtGSTU1 plays a role in the resistance of Arabidopsis to *Pst* DC3000. In contrast, chlorotic symptoms were rarely observed on the leaves of the three *TaGSTU6*-OE lines (*TaGSTU6*-OE6/13/20) compared with WT leaves (**Fig. 9A; Supplemental Fig. S22A**). RT-qPCR analysis revealed that the transcript levels of *TaGSTU6* in *TaGSTU6*-OE20 were the most elevated, followed by *TaGSTU6*-OE13 and *TaGSTU6*-OE6, which is consistent with their levels of resistance (**Fig. 9B**). Furthermore, pathogenic bacterial growth was significantly inhibited in the *TaGSTU6*-OE lines compared with the *gst* and WT plants (**Fig. 9C**). These results thus verify that *TaGSTU6* increased the resistance of Arabidopsis to *Pst* DC3000, and Arabidopsis resistance is positively correlated with *TaGSTU6* expression levels. We then examined whether overexpression of *TaCBSX3* also increases the resistance of Arabidopsis to *Pst* DC3000. Compared with WT lines, chlorotic symptoms were obvious reduced on leaves of the three *TaCBSX3*-OE lines (**Fig. 9D; Supplemental Fig. S22B**). Moreover, the anti-bacterial effects were consistent with the expression level of the *TaCBSX3*-OE lines (**Fig. 9E**). As shown in Fig. 9F, pathogenic bacterial growth was also significantly inhibited in the *TaCBSX3*-OE lines compared with WT plants. In conclusion, overexpression of *TaGSTU6* or *TaCBSX3* enhanced disease resistance to *Pst DC3000* in Arabidopsis.

**Figure 9.**
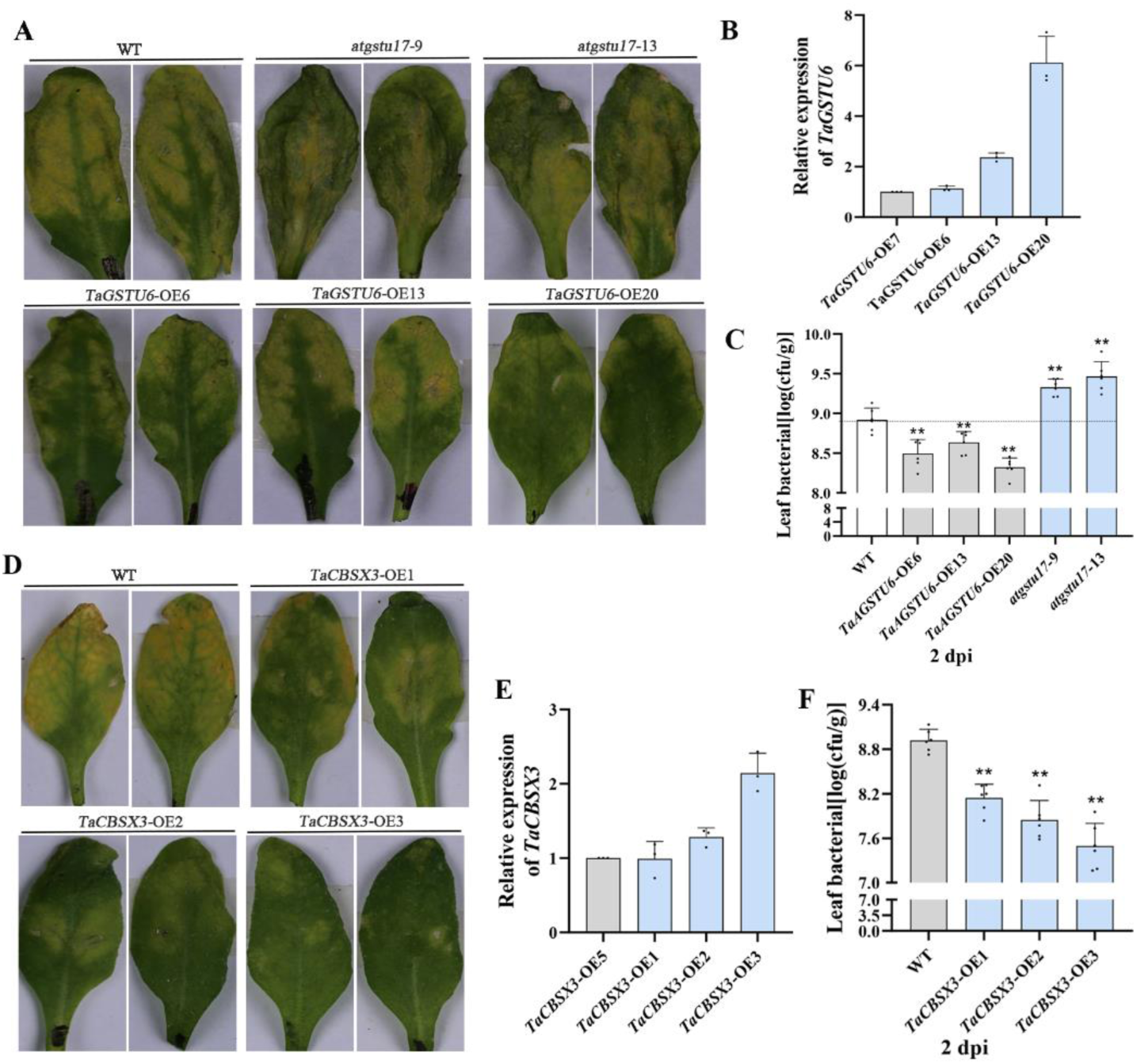
Overexpression of TaGSTU6 and TaCBSX3 promotes Arabidopsis resistance to *Pst* DC3000. A and D, Phenotypes of *TaCBSX3*-OE and *TaGSTU6*-OE overexpression lines, *gst* mutant and WT Arabidopsis at 2 dpi with *Pseudomonas syringae* pv. *tomato* (*Pst*) DC3000. *TaGSTU6* overexpression lines were more resistant than WT and *gst* mutants, and the *TaCBSX3* overexpression lines were more resistant than WT plants. B and E, Relative transcript levels of *TaGSTU6* and *TaCBSX3* in *TaGSTU6*-OE and *TaCBSX3*-OE lines by RT-qPCR. C and F, Bacterial titers (log10) of the *TaCBSX3*-OE, *TaGSTU6*-OE overexpression lines and the *gst* mutant at 2 dpi with *Pst* DC3000 (OD_600_ = 0.01). Bacterial growth is expressed as mean values of viable bacteria per gram of leaf tissue ± SD. Error bars indicate SD. Data are shown as mean ± SD (n = 6, where n indicates biological replication). Differences were assessed using Student’s t tests. \**P* < 0.05; \*\**P* < 0.01. WT, wild type.

## Discussion

Stripe rust and powdery mildew are important worldwide diseases of wheat. Transcriptome surveys demonstrated that different pathways and genes were activated in wheat leaves upon *Pst* and *Bgt* infection (Zhang et al. 2014). In this study, we have extended this work by performing quantitative proteomics to analyze the mechanisms of wheat resistance interactions with *Pst* and *Bgt*. Our results show that proteins in different functional categories are activated in response to *Pst* and *Bgt,* and that there are obvious differences and correlations between the DAPs in *Pst* and *Bgt* infection.

### Different Protein Kinases are Involved in Wheat Responses to *Pst* and *Bgt*

Mitogen-activated protein kinases (MAPKs), a group of highly conserved proteins, are key signaling enzymes involved in the regulation of various plant processes, including metabolism and abiotic and biotic defense signaling processes (Xu and Zhang, 2015; Liu et al., 2017). Accumulation of MPK3 and MPK6 is critical for stimulating full induction of plant defense during induced resistance (Gawroński et al., 2014). *TaMAPK4* was induced during the wheat-*Pst* incompatible interaction and was regulated by miR164, which indicated that MAPK signaling plays an important role in wheat defense against the rust pathogen (Wang et al., 2018). Li et al. (2017) found that the MAPK5 transcripts were reduced in *Bgt* infected susceptible wheat leaves. Therefore, they postulated that the MAPK5 may have a positive role in wheat resistant to powdery mildew. In this study, consistent with the results of Li et al. (2017), the MAPK5 protein level was significantly increased in the incompatible wheat-*Bgt* interaction, but not in the incompatible wheat-*Pst* interaction, suggesting that MAPK5 plays a role in wheat resistance to *Bgt* but not to *Pst*.

Wheat calcium-dependent protein kinases (CDPKs) are crucial sensors of calcium concentration changes that occur in response to various biotic and abiotic stressors, including H_2_O_2_, salt, drought, and *Bgt* (Li et al., 2008). Freymark et al. (2007) found that at least nine CDPK paralogs are expressed during the early phases of powdery mildew infection of barley (*Hordeum vulgare*). Similarly, in our study, CDPK2 expression was up-regulated in the wheat-*Bgt* interaction. Therefore, we speculate that MAPK5 and CDPK2 have important roles in wheat resistance to *Bgt*. In addition, other classes of protein kinases were identified in the wheat interactions with *Pst* and *Bgt* (**Supplemental Table S5**), but no protein kinase was found to be affected in both *Pst* and *Bgt* infections, suggesting that wheat utilizes different kinases in response to *Pst* and *Bgt* stresses.

### Pathogenesis-Related Proteins (PRs) Highlight Differences in Wheat Resistance to *Pst* and *Bgt*

PRs are types of proteins produced by plants under various biological or abiotic stresses and are closely related to the hypersensitive response and to systemic acquired resistance (Van Loon and Van Strien, 1999). To date, PRs have been divided into 17 families (Sels et al., 2008), and different types of PRs have different and extremely important roles in various plant stress responses. In this study, we have identified many types of disease-resistant proteins, including PR-1, PR-2, and PR-3 in the wheat-*Bgt* interaction, and PR-1, PR-2, and Wrab17 in the wheat-*Pst* interaction. These results suggest that wheat has complex disease resistance systems that elicit differences in responses to the *Pst* and *Bgt* pathogens.

Plant lipid transfer proteins (LTPs, PR-14) also take part in broad-spectrum resistance to pathogens. Several studies have found that particular plant LTPs have various functions in inhibiting the growth of different pathogens or enhancing host resistance to various pathogens (Molina et al., 1993; García-Olmedo et al., 1995; Isaac Kirubakaran et al., 2008). *TaDIR1-2*, a novel wheat ortholog of LTP, has been confirmed as a negative regulator in wheat resistance to *Pst* by modulating ROS (Ahmed et al., 2017). LTPs also participate in the abiotic stress responses. Wheat *TdLTP4* is induced by salt, drought, ABA, and JA, and overexpression of *TdLTP4* in Arabidopsis promotes plant growth under various abiotic stresses (Safi et al., 2015). We also identified some LTPs in wheat during *Pst* and *Bgt* infections, but the types of LTPs were inconsistent, and more LTPs were identified in wheat inoculated with *Bgt* than *Pst* (**Supplemental Table S5**). Therefore, different LTPs may be involved in the resistance response to *Pst* and *Bgt*. In addition, the wheat cultivar XM318 used in this study is resistant to both *Bgt* and *Pst*, and as Fig. 1, Fig. S1, S2 and Supplemental Table S1 showed, *Bgt* development in wheat is more rapid than *Pst*. Therefore, there is a possibility that the identified PR proteins may vary as a consequence of differential pathogen development times, which requires more intensive proteomics monitoring at different infection time points.

### Hormonal Responses Differ in Wheat-*Pst* and Wheat-*Bgt* Interactions

Plant hormones play important roles in plant defense responses to a variety of pathogens. SA has been demonstrated to be critical to immunity signaling processes and elevated SA can protect plants from a wide range of pathogens (Vlot et al., 2009). SA and its analogs also induce systemic acquired resistance in plants and induce expression of pathogenesis-related proteins (Kohler et al., 2002). ABA regulates abiotic stress resistance and negatively regulate plant disease resistance pathways (Asselbergh et al., 2008). The ET and JA pathways, also known as the JA/ET signaling pathway, are essential for induced systemic resistance of plant (Pieterse and Van Loon, 1999), and *TaWRKY62* positively regulates wheat high-temperature seedling resistance to *Pst* by regulating SA, ET, and JA-mediated signaling (Wang et al., 2017). These signaling pathways interact with each other through positive and negative network interactions, and ultimately, plants appear to use the most reasonable defense response combinations against specific pathogens (Flors et al., 2008).

In this study, many hormone-related proteins were identified in wheat-*Pst* and wheat-*Bgt* interactions (**Supplemental Table S5**), but few were simultaneously induced in both interactions. Among these hormone-related proteins, ABA- and SA-related proteins were activated in response to *Pst,* and SA-, JA-, and ABA-related proteins were mainly expressed in *Bgt* responses. Huai et al. (2019) found that *Pst* infection resulted in increased ABA accumulation, and our determination of endogenous hormone contents by HPLC-MS/MS further confirmed it. In wheat-*Pst* interaction, the ABA and SA contents in leaves were significantly up-regulated. In contrast, the SA and ACC levels increased significantly in *Bgt*-infected wheat leaves, but the JA concentrations were down-regulated (**Fig. 8, C and D**). These results show that hormonal responses are regulated differently in wheat-*Pst* and wheat-*Bgt* interactions. Further analyzes indicated that SA accumulation decreased significantly in *TaGSTU6-* or *TaCBSX3*-silenced leaves, whereas other hormones were almost unchanged or had minor changes after inoculation with *Bgt* (**Fig. 8, E and F**). Therefore, SA plays an important role in *TaGSTU6* and *TaCBSX3* conferring resistance to *Bgt* infection. Taken together, the ABA or SA pathway or both appear to be involved in wheat defenses against *Pst*, whereas single SA, JA, or ACC pathways or a combination of these pathways participate in defense responses against *Bgt*.

### TaGSTU6-TaCBSX3 Interaction Promotes Wheat Resistance to *Bgt*

GSTs constitute an ancient, rich class of multifunctional proteases and are encoded by a multi-gene family. A genome-wide comprehensive analyses have identified 346 *GST* genes and 87 *TaGSTU* members in common wheat (Hao et al., 2021; Wang et al, 2016). RT-qPCR assays show that the *TaGSTU6*, *TaGSTU4* and *TaGSTU7* are significantly up-regulated in the incompatible wheat-*Bgt* interaction (**Supplemental Fig. S23**). But the transcripts of *TaGSTU4* and *TaGSTU7* were lower than *TaGSTU6* during *Bgt* infection.

The previous study indicated that *Blumeria* effector BEC1054 also interacted with barley Glutathione-S-transferases (GST) to confer plant defense with ROS accumulation (Pennington et al., 2016). Li et al. (2017) reported that *TaGSTU61* was involved in oxidative burst in response to *Bgt* infection, suggesting that the high levels of antioxidants may protect wheat from oxidative damage. Therefore, we hypothesized that *TaGSTU6* is involved in wheat resistance to *Bgt* in an ROS-dependent manner.

In this study, mass spectrometry and RT-qPCR results showed that *TaGSTU6* is significantly regulated in both *Pst* and *Bgt* infections. However, the VIGS assay showed that *TaGSTU6* acts as a resistance-associated protein in the wheat response to *Bgt* infection, but not to *Pst.* In summary, *TaGSTU6* has different roles in wheat resistance to *Pst* and *Bgt*. There are six CBS domain-containing proteins predicted in the Arabidopsis genome, and these contain only a single CBS pair, but no protein domains were designated as CBSX (Ok et al., 2012). CBSXs are speculated to be novel sensor relay proteins that function in regulation of many enzymes (Seok et al., 2020). CBSX1 and CBSX2 directly regulate activation of TRXs as redox regulators in chloroplasts. Overexpression of *CBSX1* and *CBSX2* reduces ROS levels, and subsequently affects the expression of cell wall thickening genes (Yoo et al., 2011; Jung et al., 2013). Interactions of CBSX3 with Thioredoxins (Trx-o2) provide a ROS mitochondrial regulator in Arabidopsis and ROS accumulation was shown to increase markedly during *CBSX3*-overexpression (Seok et al., 2020). The interaction of the transcription factors TaNAC069 and TaCBSX3 may have positive regulatory roles in wheat resistance to *P. triticina* (Zhang et al., 2021). Mou et al. (2015) determined that *OsCBSX3* acts as a positive regulator in rice resistance to *M. oryzae* by working synergistically through SA and JA-mediated signaling pathways. OsCBSX4 was developmentally regulated by sense alterations in ionic and energy balance (Singh et al., 2012). Therefore, TaCBSX3 may directly activate Trx and regulate the production of ROS.

Our results indicated that TaCBSX3 associates with TaGSTU6 during wheat-*Bgt* incompatible combinations. Additional BMSV-VIGS and transgenic Arabidopsis inoculation assays showed that *TaCBSX3* can positively regulate resistance to *Bgt* and to the bacterial *Pst* DC3000 pathogens, respectively. Therefore, we speculate that wheat senses *Bgt* signals after infection, and that TaGSTU6 may interact with effector proteins secreted by *Bgt*. After infection, TaGSTU6 interacts with TaCBSX3 to pass signals into mitochondria, and TaCBSX3 associates with Trx to activate downstream genes that regulate ROS production and inhibit *Bgt* growth. The accumulation of ROS and SA are important signal factors in plant disease resistance responses, and are involved in regulating and inducing the expression of a series of disease resistance and systemic acquired resistance genes (Liu, 2020). Increased SA levels were detected in Arabidopsis plants with persistent H_2_O_2_ production by peroxisomes (Oa et al., 2010). Salicylic acid-mediated defense against *Pseudomonas syringae* was impaired in Arabidopsis plants with insufficient peroxidase production ROS (Mammarella et al., 2014). Thus, TaGSTU6 interacting with TaCBSX3 was involved in the regulation of reactive oxygen species concurrently, so that the plant produces disease resistance response while the plant accumulates SA signaling molecules at the pathogenic infestation site, which induces the plant to produce SAR. As a regulator of *Bgt* resistance, TaGSTU6 may participate in numerous ways to inhibit *Bgt* infection. Verification of these activities requires further studies of disease resistance mechanisms.

## Conclusions

This study reveals that wheat has different mechanisms in response to obligate biotrophic parasitic fungi, *Pst* and *Bgt*, and differences exist in hormonal responses and protein kinase activities. We also confirmed that TaGSTU6 associated with TaCBSX3 enhances wheat resistance to *Bgt*, but has little apparent effect against *Pst*. Furthermore, TaGSTU6 associations with TaCBSX3 positively regulate wheat resistance to *Bgt* through the SA signaling pathways. Our results provide new insights into the wheat resistance mechanisms and new resources for disease-resistance breeding and transgenic resistance.

## Materials and Methods

### Biological Materials, Fungal Inoculation, and Chemical Treatments

The Chinese winter wheat line XM318, developed by Shaanxi Xingmin Seed Company, shows high resistance to all current Chinese predominant *Pst* and *Bgt* races and pathotypes. The *Pst* isolate CYR32 and the *Bgt* isolate E09, which are predominant races in China (Xia et al., 2020; Wan et al., 2007), were used in this study. CYR32 and E09 were maintained and propagated on the susceptible Chinese wheat cultivars Mingxian169 (MX169) and Jingshuang16 (JS16), respectively. *N. benthamiana* and Arabidopsis (Columbia-0 background) were used in this study, and plant cultivation was performed as previously described (Huai et al., 2019).

For inoculation, XM318, MX169, and JS16 seeds were planted at the same time under identical and fungal-free conditions. When the first seedling leaves had fully expanded, half of the XM318 seedlings were brushed with *Pst*, and the other half of the XM318 seedlings were inoculated with *Bgt*. MX169 and JS16 seedlings were inoculated with *Pst* and *Bgt*, respectively, as susceptible controls. Leaves were collected at 12, 24, 36, 48, 60, 72, 84, and 96 hpi for histological observations and RT-qPCR analyses. In addition, the leaves were sampled at 0, 24, and 48 hpi for proteomics.

For chemical treatments, XM318 (incompatible) and JS16 (compatible) seedlings were treated with SA, ET, JA and ABA, as previously described (Wang et al., 2020). Leaves were sampled at 0, 0.5, 2, 6, 12, and 24 h posttreatment. For endogenous hormone accumulation, *TaGSTU6*-silenced and *TaCBSX3*-silenced leaves inoculated with *Pst* or *Bgt* (incompatible combination) were sampled at 8 and 12 hpi for HPLC-MS/MS analysis. For gene silencing assays, the wheat cultivars XM318 and “Suwon 11” BSMV-silenced leaves were inoculated with *Pst* or *Bgt* as incompatible and compatible combinations, respectively.

All of the sampled leaves were immediately frozen in liquid nitrogen and stored at - 80℃. Three different replications were performed for each treatment.

### Histological Staining

For histological observations, sampled leaves were stained with 3,3’- diaminobenzidine hydrochloride (MP Biomedicals, USA; Xiao et al., 2003) and wheat germ agglutinin (WGA; Invitrogen, USA) as described previously (Ayliffe et al., 2011). Necrotic areas were detected by auto-fluorescence of mesophyll cells in infected leaves by epi-fluorescence microscopy. For each treatment, 50 random infection sites were examined microscopically with an Olympus BX-53 microscope (Olympus, Japan). ROS areas generated by H_2_O_2_, hyphal branches, hyphal length, and areas of infection were observed and measured.

### Total Protein Extraction and TMT Labeling

For proteomics, total proteins were extracted from samples using the trichloroacetic acid /acetone method as previously described (Zhang et al., 2016). For each sample, 200 μg of proteins were digested with 4 μg solid trypsin according to a filter-aided sample preparation procedure (Wiśniewski et al., 2009). Then peptide mixtures of each sample were labeled using the TMTsixplex reagent according to the manufacturer’s instructions (Thermo Fisher Scientific). Samples were labeled with the TMT tags, and three biological repeats had three sets of TMT tags as detailed in Supplemental Table S9.

### LC-MS/MS Analysis and Proteomic Data Analysis

The Pierce High pH Reversed-Phase Fractionation Kit (Thermo Scientific) was used to separate the TMT-labeled digested samples into 15 fractions. LC-MS/MS analysis of each fraction was performed with a Q Exactive mass spectrometer (Thermo Scientific) as described by Zhang et al. (2017). Raw MS/MS data were searched using the MASCOT engine (Matrix Science, London, UK; version 2.2) embedded into Proteome Discoverer 1.4 (Thermo Scientific). After the raw data (.raw) was converted to a .mgf file, it was then run against the protein database: uniprot_Triticum_aestivum_146090_20170302.fasta (Total number of sequences: 146090, download link: http://www.uniprot.org). The following parameters were set: the peptide FDR was set to ≤ 0.01; the variable modification choice was oxidation (M) and TMT 6plex (Y); for fixed modification, Carbamidomethyl (C), TMT 6plex (N-term) and TMT 6plex (K) were selected. Other search parameters were set according to Guo et al. (2020). For protein quantification, the mock inoculation sample at 0 h was used as a reference. The quantitative protein ratios were weighted and normalized by the median ratio in Mascot. DAPs were analyzed according to the ratios of the abundance of the proteins identified and a one-sample t-test. Proteins with *p*-values (*t*-test) < 0.05 and fold change ratios ≥1.3 or ≤0.77 were considered to be significantly upregulated or downregulated, respectively.

### Bioinformatics Analysis

Functional annotations of the identified proteins were conducted using the Blast2GO program against the UniProtKB database. Level 2 GO analyses of multi-group data were performed with OmicShare tools, a free online platform for data analysis (http://www.omicshare.com/tools). The expression levels of all DAPs were converted into log2 fold changes. Then, R language was used to perform heatmap analyses of all identified proteins and DAPs. DAPs were classified into different functional categories with the COG database (http://eggnogdb.embl.de/#/app/emapper?jobname=MM_FnYQKJ). Analysis of plant visualization pathways was performed using MapMan software. KAAS (KEGG Automatic Annotation Server) was used to retrieve their Kos, which were subsequently mapped to KEGG pathways, and TBtools software was used to draw the KEGG enrichment chart.

### Total RNA Extraction and RT-qPCR

With reference to the Plant RNA Reagent instructions (TransGen Biotech, China), total RNAs were isolated and reverse transcribed into cDNA following the manufacturer’s directions (Thermo, USA), and genomic DNA contaminants were removed by DNase I treatment. Specific primers (Supplemental Table S10) were designed using Primer Premier 5.0 software to assess the transcriptional expression levels of selected protein genes by RT-qPCR. The wheat elongation factor *TaEF-1α* was chosen as the internal control gene. All RT-qPCR was performed according to the manufacturer’s instructions (TransGen Biotech, China). Cycle threshold values were generated using the QuantStudio 5 Real-Time PCR System (Applied Biosystems, USA) to quantify relative gene expressions by use of the comparative 2^−ΔΔCt^ method (Livak and Schmittgen, 2001). All PCR analyses were repeated three times.

### Sequence Analysis of *TaGSTU6* and *TaCBSX3*

*TaGSTU6* and *TaCBSX3* alleles from chromosomes were obtained from the wheat UGRI genome database (https://urgi.versailles.inra.fr/). The NCBI conserved domain database (https://www.ncbi.nlm.nih.gov/cdd/) was used to identify domains. Phylogenetic trees were constructed with MEGA7.0 software. Physicochemical properties of proteins were predicted using the “ProtParam” tool of “ExPASy” (http://www.expasy.org).

### BSMV-Mediated Gene Silencing

The *TaGSTU6* and *TaCBSX3* genes were silenced using the BSMV-VIGS system. Two pairs of special gene fragments (*TaCBSX3*: 255 bp and 362 bp; *TaGSTU6*: 199 bp and 282 bp) were generated and cloned into BSMV as previously described (Holzberg et al., 2002). The infectious BSMV RNAs were transcribed in vitro from linearized plasmids γ-*TaPDS*, γ-*TaCBS*, γ-*TaGSTU6*, γ, α, and β (Petty et al., 1989) using the RiboMAX Large Scale RNA Production System-T7 kit (Promega, USA) and the Ribo m7G Cap Analog (Promega). Each group of mixed BSMV RNA was inoculated onto the second leaf of wheat seedlings at the two-leaf stage as described previously (Scofield et al., 2005). The seedlings were maintained in a growth chamber at 25℃ and examined for symptoms. BSMV:*TaPDS* acted as the positive control, BSMV:γ was used as the viral control, and mock plants were inoculated with FES buffer as the negative control. At 10-12 dpi with BSMV, at the four-leaf stage of seedlings, the third and fourth leaves of each plant were inoculated with *Pst* or *Bgt*, and leaves were sampled at 0, 24, 48, 96, and 120 hpi to determine the silencing efficiencies and histological observations. The stripe rust and powdery mildew phenotypes were evaluated at 16 and 12 dpi. All primers used are listed in Supplemental Table S10, and the experiments were repeated three times.

### Y2H Assay

The cDNA libraries for the Y2H assay were constructed using the Gateway system (Invitrogen, USA) by OE Biotech (China). The cDNAs were constructed from total RNAs extracted from *Bgt* -infected XM318 leaves. The pGADT7 (AD)-cDNA library vector was used as the prey. The bait vectors were generated by inserting *TaGSTU6* DNA fragments into pGBKT7 (BD) plasmids. The bait and prey vectors were co-transformed into yeast strain Y2H Gold according to the Yeast Protocols Handbook (Clontech), diluted on SD/-Leu-Trp/AbA for selection of positive colonies, which were transferred to SD/-Leu-Trp-His-Ade/X-α-Gal/AbA for further selection. To confirm *TaGSTU6* interaction targets, full-length cDNAs of the candidate target of interest were cloned into pGADT7 vectors and transformed into the yeast strain Y2H Gold, containing the bait vector, and grown on SD/-Leu-Trp-His-Ade/X-α-Gal/AbA.

### BiFC Assay

For the BiFC assays, *TaGSTU6* cDNAs were cloned into pSPYNE to make nEYFP-*TaGSTU6* constructs, and the cDNA from *TaCBSX3* was ligated into pSPYCE to form cEYFP-*TaCBSX3*. The constructed plasmids were transformed into *A. tumefaciens* GV3101. The *Agrobacterium* strains were infiltrated into the leaves of 4-week-old *N. benthamiana* in different combinations. Two days after inoculation, images were observed via Olympus BX-53 microscope (Japan). The H2B-mcherry fusion protein was used as a nuclear marker.

### Firefly Luciferase Complementation Imaging (LCI) assays

For LCI assays, *TaGSTU6* and *TaCBSX3* were cloned into pCAMBIA1300-NLuc and pCAMBIA1300-CLuc to confirm TaGSTU6 interactions with TaCBSX3. Primers used for vector construction are shown in Supplemental Table S10. LCI assays were performed in *N. benthamiana* leaves as previously described (Chen et al., 2008). Three independent replicates were performed.

### Co-IP and Western Blot Analysis

For preparation of Co-IP assays, protein samples, and colocalization of TaGSTU6 and TaCBSX3, full-length *TaGSTU6* and *TaCBSX3* cDNAs were amplified to construct pCAMBIA1302-TaGSTU6-GFP and pYBA1132-TaCBSX3-mCherry plasmids. The primers used are listed in Supplemental Table S10. *Agrobacterium* strains containing TaGSTU6-GFP and TaCBSX3-mCherry plasmids were transformed into *N. benthamiana* leaves at an OD_600_ of 0.5. At 48 h after agroinfiltration, fluorescence signals were observed with an Olympus BX-53 microscope (Olympus, Japan), and *N. benthamiana* total proteins were extracted with a native lysis buffer (Solarbio).

For Co-IP assays, 15 μL of GFP-Trap agarose beads (Chromotek, Germany) were incubated with 2 mL of crude protein at 4°C for 3 h and collected as described previously (Yang et al., 2019).

For western blot analyses, SDS-PAGE, electroblotting, and immunodetection analysis were carried out as described previously (Yang et al., 2020). Precipitated proteins were detected with an anti-GFP antibody (#A02020; Abbkine), an anti-mCherry antibody (#A02080; Abbkine), and a goat anti-mouse secondary antibody (ab6789; Abcam). Protein bands were visualized with a western ECL substrate kit (Bio-Rad).

### Arabidopsis Transformation and Inoculation

The recombinant plasmid pCAMBIA2300-TaGSTU6 and pCAMBIA2300-TaCBSX3 were constructed, transferred into *Agrobacterium tumefaciens* strain EHA105, and subsequently transformed into Arabidopsis ecotype Columbia-0 as described (Zhang et al., 2006). More than 30 independent transgenic lines were screened based on resistance to the antibiotics G418 and kanamycin. The increased transcript levels of *TaGSTU6* and *TaCBSX3* were verified by RT-qPCR. Fully expanded leaves of 4-week-old transgenic Arabidopsis were infected with suspensions of *Pst* DC3000 cells (OD_600_=0.01) in 10 mM MgCl_2._ Bacterial growth in plant leaves was determined at two dpi. Inoculation and conidiophore counting methods were as described by Xiao et al. (2005).

### Accession numbers

All mass spectrometry proteomics data in this study have been deposited in the Integrated Proteome Resources database (iProX), a member of the ProteomeXchange Consortium, and can be browsed at the iProX webpage with the accession number IDX0001436000. Sequence data from this article can be found through the NCBI database (http://www.ncbi.nlm.nih.gov/) with the accession number *TaGSTU6* (XM_044472374.1), *TaCBSX3* (XM_020317884.1) and *AtGSTU17* (AY091102.1).

## Acknowledgements

We thank Prof. Andrew Otis Jackson, Dr. Qiaojun Jin, and Charlesworth Author Services for critically editing the manuscript.

## Supplemental Data

**Supplemental Figure S1.** Histological changes in *Pst* growth, H_2_O_2_ accumulation, and necrotic cell areas in wheat XM318 and MX169 cultivars.

**Supplemental Figure S2.** Histological changes in *Bgt* growth, H_2_O_2_ accumulation, and necrotic cell areas in wheat XM318 and JS16 cultivars.

**Supplemental Figure S3.** The abscissa shows scoring of the MASCOT peptide fragment and is the relative molecular weight of the identified protein.

**Supplemental Figure S4.** Protein profiles identified by TMT in *Pst* and *Bgt* infected wheat.

**Supplemental Figure S5.** Classification and regulation of DAPs.

**Supplemental Figure S6.** GO annotation of HRPs in the TMT data by Blast2GO software.

**Supplemental Figure S7.** Analysis of pathways for wheat interactions with *Pst* and *Bgt* at different infection times.

**Supplemental Figure S8.** KEGG pathway enrichment analysis of DAPs.

**Supplemental Figure S9.** Twelve DAPs genes were selected and RT-qPCR analyses performed.

**Supplemental Figure S10.** *TaGSTU6* identification and transcription profiles.

**Supplemental Figure S11.** Functional analysis of *TaGSTU6* in wheat resistance to *Pst* after BSMV-induced gene silencing.

**Supplemental Figure S12.** Fungal growth rates in incompatible interactions of *TaGSTU6*-knockdown plants.

**Supplemental Figure S13.** Functional assessment of *TaGST* roles in compatible wheat-*Bgt* interactions after BSMV gene silencing.

**Supplemental Figure S14.** Epifluorescence observations of *Bgt* growth in wheat in compatible interactions after inoculation with BSMV.

**Supplemental Figure S15.** Interactions of TaGSTU6 with TaCBSX3 in yeast two-hybrid assays.

**Supplemental Figure S16.** *TaCBSX3* Expression profiles in response to *Bgt* and *Pst* infection.

**Supplemental Figure S17.** Functional analysis of *TaCBSX3* in *Pst* resistant wheat after BSMV-induced gene silencing.

**Supplemental Figure S18.** Fungal growth rates in incompatible interactions of *TaCBSX3*-knockdown plants.

**Supplemental Figure S19.** Functional assessment of the role of *TaCBSX3* determined by BSMV-mediated gene silencing in compatible wheat-*Bgt* interactions.

**Supplemental Figure S20.** Epifluorescence observations of *Bgt* compatible growth in wheat after BSMV inoculation.

**Supplemental Figure S21.** *TaGSTU6* and *TaCBSX3* induction after exogenous hormone treatments.

**Supplemental Figure S22.** Bacterial defense phenotypes in *TaCBSX3*-OE and *TaGSTU6*-OE transformed Arabidopsis.

**Supplemental Figure S23.** Expression profiles of some *TaGSTU*s.

**Supplemental Table S1-1.** Numbers of HMC, hyphal branches, linear lengths, and infection foci in *Pst* infected wheat.

**Supplemental Table S1-2.** Numbers of haustoria, hyphal branches, linear length, and areas of infection per infection site in the *Bgt* wheat-interaction.

**Supplemental Table S2.** Percentage of infection sites of mesophyll cells exhibiting H_2_O_2_ accumulation and necrotic cells after different infection interactions.

**Supplemental Table S3.** Results of protein identification statistics.

**Supplemental Table S4.** DAPs regulated during *Pst* and *Bgt* infections.

**Supplemental Table S5.** Differentially expressed protein groups in wheat infected with *Pst* or *Bgt*.

**Supplemental Table S6.** Regulation multiple of DAPs selected for RT-qPCR.

**Supplemental Table S7.** *TaGSTU6* and T*aCBSX3* transcription profiles and protein expression level in response to *Bgt* and *Pst* infection.

**Supplemental Table S8.** Description of potential TaGSTU6 targeted by the Y2H system.

**Supplemental Table S9.** Sample tag information in TMT analysis.

**Supplemental Table S10.** Primers used for PCR and plasmid construction.

**Supplemental file S1.** All identified proteins in the wheat-*Pst* and wheat-*Bgt* interaction.

**Supplemental file S2.** Details of differentially expressed proteins in the wheat-*Pst* interaction.

**Supplementary file S3.** Details of differentially expressed proteins in the wheat-*Bgt* interaction.

**Supplementary file S4.** Details of differentially expressed proteins by comparing wheat-*Pst* and wheat-*Bgt* interactions at 24 hpi and 48 hpi.

**Supplementary file S5.** Details of differentially expressed proteins in wheat interactions with both *Pst* and *Bgt*.

**Supplementary file S6.** Lists of proteins in different COG categories.

